# Crystal structure of the GluK1 ligand-binding domain with kainate and the full-spanning positive allosteric modulator BPAM538

**DOI:** 10.1101/2023.11.02.565302

**Authors:** Yasmin Bay, Stine M. Fransen, Darryl S. Pickering, Karla Frydenvang, Pierre Francotte, Bernard Pirotte, Anders Skov Kristensen, Jette Sandholm Kastrup

**Affiliations:** Department of Drug Design and Pharmacology, Faculty of Health and Medical Sciences, University of Copenhagen, DK-2100 Copenhagen, Denmark; Department of Medicinal Chemistry, Center for Interdisciplinary Research on Medicines (CIRM), University of Liège, Liège, Belgium

**Keywords:** ray crystallography, ligand-binding domain of GluK1, positive allosteric modulator, kainate, binding mode

## Abstract

Kainate receptors play an important role in the central nervous system by mediating postsynaptic excitatory neurotransmission and modulating the release of the inhibitory neurotransmitter GABA through a presynaptic mechanism. To date, only three structures of the ligand-binding domain (LBD) of the kainate receptor subunit GluK1 in complex with positive allosteric modulators have been determined by X-ray crystallography, all belonging to class II modulators. Here, we report a high-resolution structure of GluK1-LBD in complex with kainate and BPAM538, which belongs to the full-spanning class III. One BPAM538 molecule binds at the GluK1 dimer interface, thereby occupying two allosteric binding sites simultaneously. BPAM538 stabilizes the active receptor conformation with only minor conformational changes being introduced to the receptor. Using a calcium-sensitive fluorescence-based assay, a 5-fold potentiation of the kainate response (100 μM) was observed in the presence of 100 μM BPAM538, whereas no potentiation was observed at GluK2.

**Highlights:**

- 1.9 Å structure of the kainate receptor GluK1-LBD in complex with kainate and BPAM538
- The positive allosteric modulator BPAM538 occupies two binding sites
- The binding mode is similar to class III modulators described for AMPA receptors
- BPAM538 prefers GluK1 over GluK2

## 1. Introduction

The ionotropic glutamate receptors (iGluRs) are involved in most of the fast excitatory synaptic transmission in the central nervous system. The iGluRs are categorized into four receptor subfamilies: AMPA, kainate, NMDA, and delta receptors (Hansen et al., 2021). This class of receptors is important for learning and memory formation but is also entangled in the development of neurological diseases. The kainate receptors are believed to mediate both a postsynaptic excitatory response and a modulatory presynaptic response leading to the release of the inhibitory neurotransmitter GABA (Lerma et al., 2013). Kainate receptors have also been associated with neurological diseases such as Alzheimeŕs disease, epilepsy, schizophrenia, and depression (Lerma et al., 2013).

The kainate receptors comprise five subunits, termed GluK1-5. While the subunits GluK1-3 may form functional homo- and hetero-tetrameric receptors, subunits GluK4-5 can only form functional receptors with GluK1-3 (Contractor et al., 2011). The topology of a single subunit is composed of four distinct domains that share common structural features with the other iGluR subfamilies (Sobolevsky et al., 2009) (Fig. 1A). Among the iGluRs, the physiological role of kainate receptors is less well understood and tool compounds with subtype selectivity are sparse. Understanding the molecular function of kainate receptors may help unravelling their potential as targets for treatment of neurological diseases by the discovery of selective tool compounds.

**Fig. 1.**
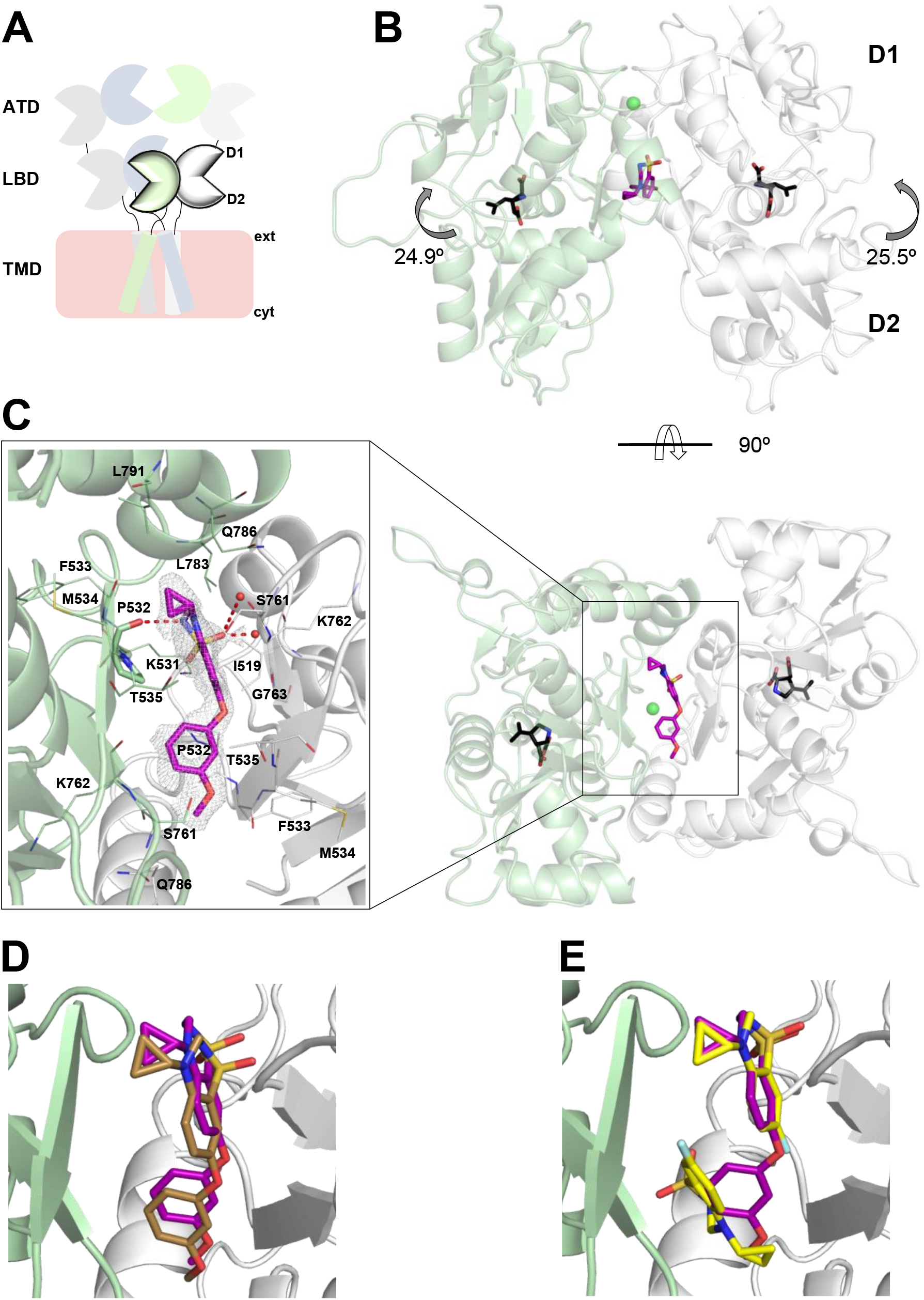
Crystal structure of GluK1-LBD in complex with the agonist kainate and the positive allosteric modulator BPAM538. **A**. Schematic drawing of a full-length kainate receptor. From the top: the amino-terminal domain (ATD) and underneath the ligand-binding domain (LBD) harboring the binding site of L-glutamate and other ligands. The LBD resembles a clamshell structure with an upper lobe D1 and a lower lobe D2. The transmembrane domain (TMD, simplified) is embedded in the membrane and connected to a carboxy-terminal domain (CTD; not shown). **B.** Dimer views: side (top panel) and bottom (bottom panel) of the GluK1-LBD structure with kainate and BPAM538. Chains A (pale green) and B (grey) are shown in in cartoon representation. Kainate and BPAM538 are shown in sticks representation with carbon atoms colored black and purple, respectively. **C.** Close-up on the lower part of the dimer interface where one BPAM538 molecule binds, presented with the 2F_o_-F_c_ omit electron density map contoured at 1 sigma and carved at 1.9 Å. Residues within 4 Å of BPAM538 are shown as lines, and water molecules as red spheres. Hydrogen bonding interactions (<3.5 Å) between BPAM538 and the residue P535 or water molecules are highlighted as red stippled lines. Oxygen atoms are red, nitrogen atoms blue, and sulfur atoms yellow. **D.** Superimposition of GluK1-LBD (kainate and BPAM538) and GluA2-LBD (L-glutamate and BPAM538; PDB code: 5OEW) structures on lobe D1 residues. GluK1-LBD is shown in cartoon representation, whereas GluA2-LBD has been left out for clarity. BPAM538 is seen to bind in a very similar manner in GluK1 (magenta) and GluA2 (brown). **E.** Superimposition of GluK1-LBD (kainate and BPAM538) and GluK1-LBD (kainate and BPAM344; PDB code: 5MFQ) structures on lobe D1 residues. BPAM538 is seen to overlay with one of the two BPAM344 molecules (represented in yellow). Only the protein chain of the structure with BPAM538 is shown.

Allosteric modulators are generally thought to be better tolerated as they modify existing levels and patterns of receptor activation rather than constitutively blocking or over-activating all receptors. Additionally, allosteric modulators have a higher potential for receptor subtype selectivity as they often target less conserved regions than the orthosteric binding site (Traynelis et al., 2010). Whereas many small molecule positive allosteric modulators (PAMs) of AMPA receptors exist, only a few PAMs for kainate receptors have been reported (Larsen et al., 2017; Puja et al., 2022). These PAMs belong to class II which bind with two molecules at the dimer interface between two ligand-binding domains (LBDs) within the receptor (Larsen et al., 2017; Frydenvang et al., 2022). PAMs exert their effect by either decreasing desensitization and/or deactivation of the receptor, thereby enhancing receptor activation by L-glutamate through binding to the allosteric site (Armstrong et al., 2000).

4-Cyclopropyl-7-(3-methoxyphenoxy)-3,4-dihydro-2*H*-1,2,4-benzothiadiazine 1,1-dioxide (BPAM538; Fig. 2) was previously reported to be a highly potent PAM at AMPA receptors with nanomolar potency (EC_50_ = 2 nM) (Goffin et al., 2018). The structure of BPAM538 in complex with the AMPA receptor GluA2 ligand-binding domain showed that BPAM538 belongs to the full-spanning class III of PAMS that occupy two allosteric binding sites per bound molecule (Frydenvang et al., 2022). In this study, we show that BPAM538 is also capable of modulating the kainate receptor GluK1 and report a 1.9 Å resolution X-ray structure of BPAM538 bound to the ligand-binding domain of the kainate receptor GluK1 with kainate.

**Fig. 2.**
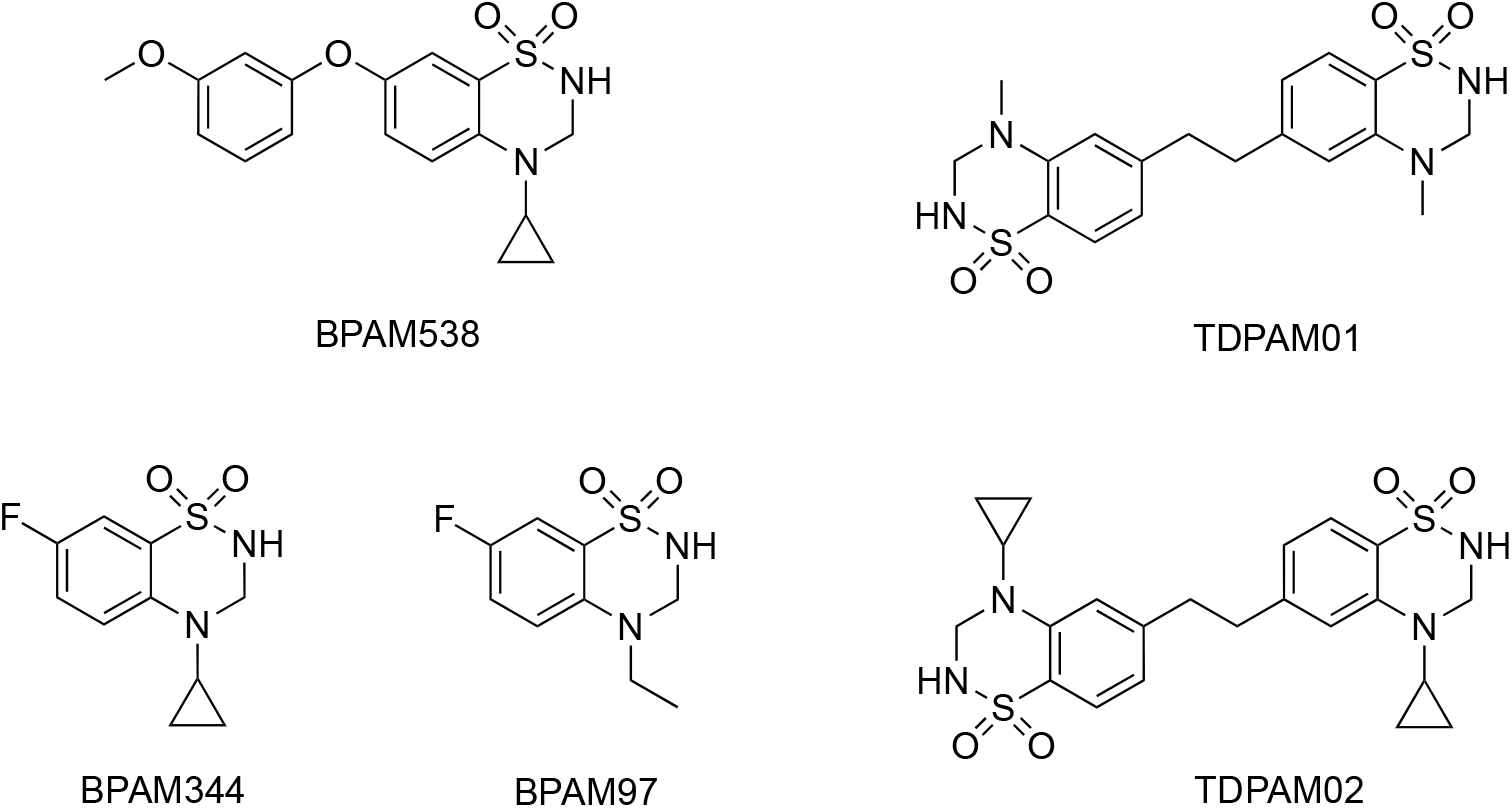
Chemical structure of the positive allosteric modulators.

## 2. Material and methods

### Structure determination

The LBD of rat GluK1 (GluK1-LBD) was expressed and purified as previously described (Naur et al., 2005). Following purification, the protein was dissolved in 10 mM HEPES pH 7.0, 20 mM NaCl, and 1 mM EDTA. BPAM538 was synthesized as previously described (Goffin et al., 2018). A final GluK1-LBD concentration of 7.5 mg/mL was used in the presence of 12.7 mM kainate saturated with BPAM538. The protein solution was equilibrated at 6 °C for at least 24 hours under rotation before setting up drops employing the hanging-drop vapor diffusion technique. A quantity of 500 µL of reservoir was added to each well in a 24-well VDX with sealant (Hampton Research). Each drop consisted of 1 µL of protein solution and 1 µL of reservoir solution (15.2% PEG4000, 0.3 M lithium sulfate, and 0.1 M sodium acetate pH 5.5). Crystals harvested for data collection were briefly submerged into the reservoir solution supplemented with 10-20% glycerol for cryoprotection and immediately flash-cooled in liquid nitrogen before storage.

X-ray diffraction data was collected at the BioMAX beamline, MAX IV Laboratory in Lund, Sweden (Ursby et al., 2020), and data was processed using XDS (Kabsch et al., 2010) and Scala (Evans et al., 2006) in the CCP4 suite of programs (Winn et al., 2011). The structure was solved by molecular replacement in PHASER (McCoy et al., 2007) using the structure of GluK1-LBD in complex with kainate (PDB code: 4E0X) (Venskutonyte et al., 2012). Initial model building was performed using AUTOBUILD (Terwilliger et al., 2008) in PHENIX (Liebschner et al., 2019). During iterative rounds of model building in COOT (Emsley et al., 2010) and refinement in PHENIX, BPAM538, kainate, glycerol, ions, and water molecules were gradually built into the structure. The ligand coordinate file of BPAM538 was generated in MAESTRO (Maestro release 2021-3; Schrödinger, LLC, New York, NY, 2021). Domain closures were calculated relative to a GluK1-LBD structure with an antagonist (PDB code: 3S2V, chain B) (Venskutonyte et al., 2011) employing the DynDom server (Hayward et al., 1998). PyMOL (The PyMOL Molecular Graphics System, Version 2.5.4 Schrödinger, LLC) was used to prepare all structure figures.

### Cell culturing and PAM activity assay

Grip Tite^TM^ Human Embryonic Kidney (GT HEK293) (Invitrogen) cells were grown as previously described (Chałupnik et al. 2022, 2023). For testing of PAM potentiation of agonist activity at GluK1(*Q*)_b_ and GluK2(*VCQ*)_a_, a Ca^2+^-sensitive fluorescence-based assay were used. Briefly, in the absence and presence of PAMs, agonist-induced increases in fluorescence intensity from an intracellular Ca^2+^-sensitive dye (Fluo8, AAT Bioquest) were measured in GluK1 and GluK2 expressing cells cultured in 96-well plates using a FlexStation I plate reader (Molecular Devices).

All PAMs were dissolved in DMSO and hereafter added to the assay buffer (100 mM choline chloride, 10 mM sodium chloride, 5 mM potassium chloride, 1 mM magnesium chloride, 20 mM calcium chloride, 10 mM HEPES, pH 7.4) on the day of the experiment. It was verified that DMSO concentrations up to 5% only had minimal effect on the agonist-induced response in GT HEK293 cells (data not shown). Attempts to improve the solubility of BPAM538 included the addition of DMSO (before and after the addition of BPAM538), pre-heating of BPAM538 stock solution for up to 45 min, assay buffer or compound solution (at least 15 min at 55°C) as well as pre-incubation of BPAM538 in assay buffer added to the cells prior to recording. However, these modifications still did not allow the determination of potency at GluK1.

Recordings of Ca^2+^ fluorescence were analyzed using SoftMax Pro software (version 5.4, Molecular Devices). Peak fluorescence response was calculated as the difference between baseline emission before agonist addition and the maximum emission after agonist addition. SigmaPlot was used to construct bar histograms shown as the mean response relative to BPAM344 (100%) ± SEM from at least four independent experiments containing eight repetitions of 100 µM PAM per plate. Statistical analysis has been used to estimate the P value between application of kainate and co-application with the tested PAMs using the Student’s t-test. A statistical difference is indicted with ** (P = <0.001) and none abbreviated as ns.

## 3. Results and Discussion

### 3.1 X-ray structure of GluK1-LBD in complex with kainate and BPAM538

In crystals, GluK1-LBD forms a dimer in the presence of kainate (Plested et al., 2008; Venskutonyte et al., 2012), creating a dimer interface with two sites for allosteric modulators. Therefore, crystallization of BPAM538 in complex with GluK1-LBD was carried out in the presence of kainate. One GluK1-LBD dimer (chain A and chain B) was observed in the asymmetric unit of the crystal, with kainate bound at the orthosteric binding site in both subunits of the dimer (Fig. 1B). Kainate displays the same binding mode as previously observed (Plested et al., 2008; Venskutonyte et al., 2012). One chloride ion was seen at the dimer interface in agreement with previous structures of kainate receptors (Plested et al., 2007) (Fig 1B).

BPAM538 was observed to bind at the dimer interface with a single molecule occupying both allosteric binding sites (Fig. 1B-E). Compared to the structure without BPAM538, the modulator seems to displace five binding-site water molecules. This defines BPAM538 as a class III PAM at kainate receptors also, as observed for AMPA receptors (Frydenvang et al., 2022). In comparison, BPAM344, belonging to class II, binds with one molecule occupying each of the two pseudo-symmetric allosteric binding sites (Fig. 1E). Residues within 4 Å of the BPAM538 molecule include K531, P532, F533, M534, T535, S761, K762, L783, Q786, and L791 from chain A and can make van der Waals interactions (Fig. 1C). Nearly, the same residues are found within 4 Å of the modulator in chain B except for K531, L791, and L783. However, additional residues from chain B appear which include I519 and G763 (Fig. 1C). Among these residues, the proline in position 532 from chain A is the only residue creating a hydrogen bond (2.8 Å) to the sulfonamide nitrogen atom of the modulator. Additionally, two water molecules in close vicinity of the modulator also make direct hydrogen bonds (2.0-3.3 Å) to one of the sulfonamide oxygen atoms in BPAM538. Five water molecules and a glycerol molecule are found close to the modulator but were not observed to generate any direct or polar contact.

Agonists are known to introduce domain closure in the LBD compared to the resting, unbound apo state of the receptor (Armstrong et al. 2000). As an apo structure is not available for GluK1, domain closures have previously been determined relative to an antagonist-bound structure (PDB code: 3S2V, chain B; Venskutonyte et al., 2011). The domain closure induced by kainate in GluK1-LBD was reported to be 25–28° (Møllerud et al., 2017). In the presence of BPAM538, the domain closure as calculated in DynDom is 24.9° and 25.5° in chains A and B, respectively (Fig.1B). Thus, domain movements and agonist binding seem to be unaffected upon binding of BPAM538. In line with this, the distances between P532, P667, and R775 of chain A and B located at the interface of the dimer inside the allosteric binding site, the bottom of lobe D2, and the top of lobe D1, respectively, were calculated. Very similar distances were measured in the crystal structure of GluK1-LBD with kainate (PDB code: 4E0X) and upon binding of BPAM538 (33.4 Å vs 32.9 Å, 9.2 Å vs 8.9 Å, and 13.3 Å vs 13.4 Å). Therefore, BPAM538 is likely to diminish desensitization of GluK1 by stabilizing the dimer interface and not by affecting the domain closure.

Superimposing the X-ray structure of GluA2-LBD complexed with L-glutamate and BPAM538 (PDB code: 5OEW; Goffin et al., 2018) onto the GluK1-LBD structure with kainate and BPAM538 gave an RMSD value of 0.62 Å when imposed on lobe D1.

BPAM538 also binds in a very similar manner in GluK1 and GluA2 (Fig. 1D-E). Similar interactions within 4 Å of BPAM538 and orientation of residues were observed between the AMPA and kainate receptor family. Of note, only two residues vary inside the binding region of BPAM538 (defined as residues within 4 Å of the modulator): T535 in GluK1 that corresponds to GluA2 S518 occurring in two conformations and Q786 in GluK1, which is N775 in GluA2. None of these residues form direct interaction with the modulator.

In comparison, a RMSD value of 0.12 Å for GluK1-LBD with kainate and BPAM538 aligned with the X-ray structure of GluK1-LBD determined with kainate and BPAM344 on lobe D1 residues (PDB code: 5MFQ; Larsen et al., 2017) was estimated, indicating high similarity between the two structures. Taking a closer look at the binding pocket and the binding mode of the two modulators, residues located within 4 Å of the modulators adopt the same orientation and BPAM538 overlaps well with one of the two BPAM344 molecules (Fig. 1E). Noteworthy, a glycerol molecule located in proximity of BPAM538 seems to occupy the space where the second BPAM344 molecule is found; yet is not able to interact with the modulator (not shown).

In conclusion, the binding mode and interactions of BPAM538 at GluK1 correlate well with what has previously been observed at GluA2.

### 3.2 Functional screen of PAMs at kainate receptors

The observation of that most PAMs work by occupying two indetical binding sites in the LBD has stimulated the design and synthesis of dimeric forms (Frydenvang et al., 2022). TDPAM02, belonging to class III, is one such example and is currently the most potent PAM at AMPA receptors known to date (Drapier et al., 2018).

We screened three class III PAMs (TDPAM01, TDPAM02, and BPAM538; Fig. 2) as well as two class II PAMs (BPAM97 and BPAM344, Fig. 2) for activity at GluK1 and GluK2. The ability of the PAMs to potentiate receptor response to 100 µM kainate was first evaluated in a single concentration of 100 µM at GluK1 using a calcium-sensitive fluorescence-based assay (Materials and Methods). Specifically, kainate activation of kainate receptors leads to an influx of Ca^2+^, which increases intracellular Ca^2+^ concentration, which can be detected using an intracellular calcium-sensitive fluorescent dye. The ability of PAMs to potentiate kainate-evoked Ca^2+^ increase was tested at a concentration of 100 µM. At this concentration, none of the tested PAMs appeared to be a better potentiator than the reference class II kainate receptor PAM BPAM344; however, BPAM97 within the same class II PAMs produced nearly similar response (87% ± 3.1%) as BPAM344 (100% ± 2.2%). As for the PAMs belonging to class III PAMs, only BPAM538 showed a modulatory effect at GluK1, potentiating the kainate response 5-fold (82% ± 5.3%) compared to (16% ± 1.5%) without PAM. TDPAM01 and TDPAM02 co-applied with kainate did not modulate GluK1 (Fig. 3A). To address KAR subtype preferences of the PAMs, BPAM538, TDPAM02, BPAM97, and BPAM344 were tested in the same manner at GluK2, which showed that only BPAM344 was observed to potentiate the response of kainate (Fig. 3B).

**Fig. 3.**
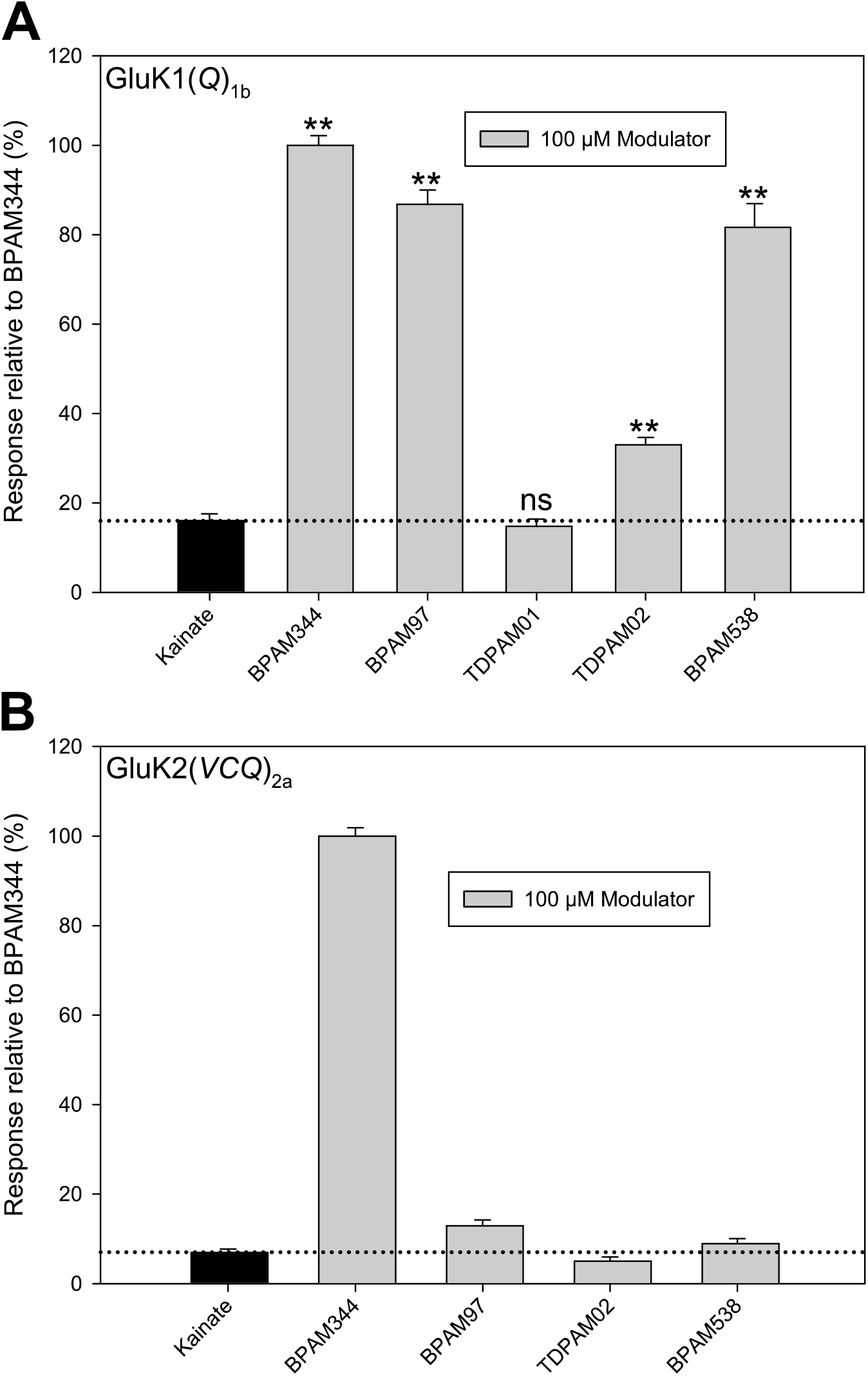
Calcium-sensitive fluorescence-based assay. **A.** Bar plots for modulation of GluK1(*Q*)_b_, representing the recorded data normalized to the BPAM344 response when co-applied with the agonist kainate (100 µM; black). BPAM344, BPAM97, TDPAM01, TDPAM02, and BPAM538 co-applied (100 µM). Response values are relative to the BPAM344 response. The bars presented are means ± SEM of eight replicate wells. The data is a representation of at least four repeated experiments. in cartoon representation **B.** Bar plots for modulation of GluK2(*VCQ*)_a_, representing the recorded data normalized to the BPAM344 response when co-applied with the agonist kainate (100 µM; black) and co-application with either BPAM97, BPAM344, TDPAM02 or BPAM538. ** = P<0.001, ns = not significant.

In summary, BPAM538 and BPAM97 showed a preference for the GluK1 receptor subtype, and characterization of the functional profile (EC_50_) was attempted using the calcium-sensitive fluorescence-based assay. However, due to the poor solubility of BPAM538 (Goffin et al., 2018) in the assay buffer, it was not possible to determine the EC_50_ value at GluK1.

### 3.3 Concluding remarks

In this study, we have determined the structure of the LBD of GluK1 in complex with kainate and BPAM538. We demonstrate that a small molecule PAM belonging to class III, manifested as one of the most potent AMPA receptor PAMs, can also modulate GluK1 expressed in HEK293 cells. On the other hand, no potentiation of GluK2 was observed, indicating that it should be possible to achieve selective modulation of kainate receptor subunits. The structural and functional insight into the new PAM class at kainate receptors provide improved basis for future design of PAMs with selectivity at kainate receptors.

**Table 1.**
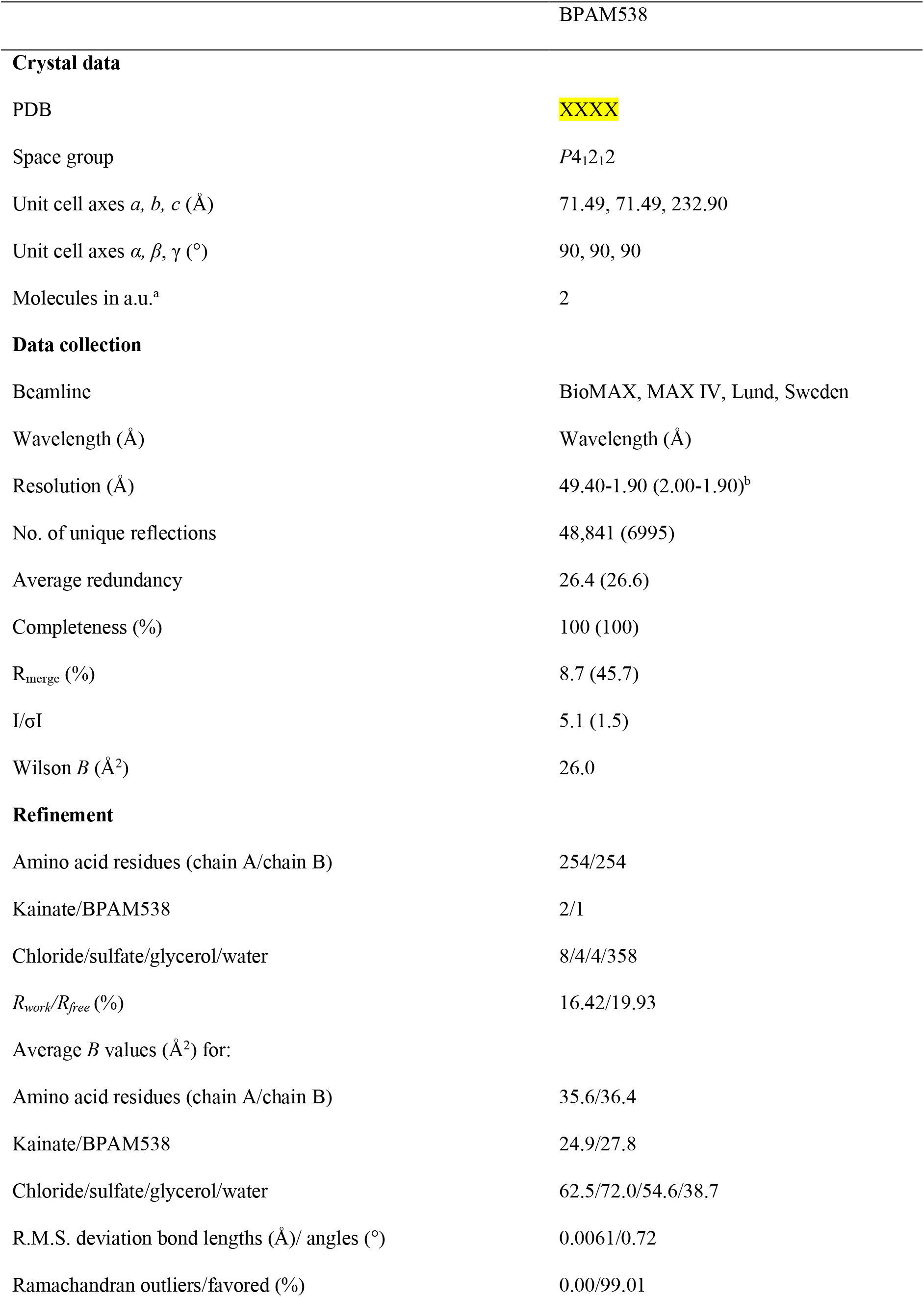
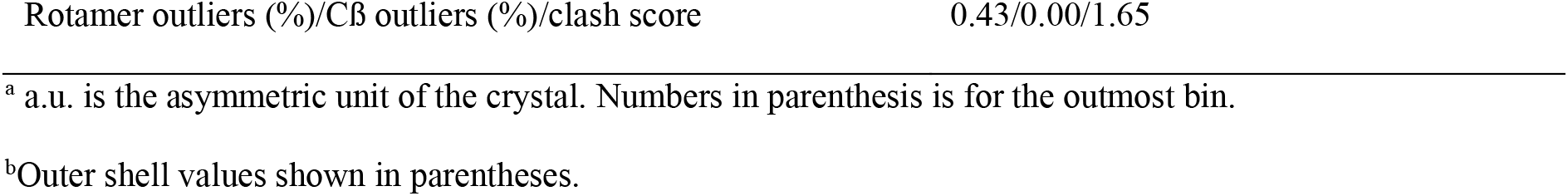
Crystal data, data collection, and refinement statistics of GluK1-LBD in complex with kainate and BPAM538.

## Acknowledgements

MAX-lab, Lund, Sweden is thanked for providing beamtime and technical assistance. Heidi Peterson is thanked for producing the GluK1-LBD protein. The Lundbeck foundation (S.M.F, J.S.K.), the Independent Research Fund Denmark – Medical Sciences (Y.B., J.S.K.) and Danscatt (Y.B., S.M., K.F., J.S.K.) are acknowledged for financial support. We acknowledge the MAX IV Laboratory for time on Beamline 911-3 under Proposal MX20130012.

## Authorship Contributions

**Yasmin Bay**: Conceptualization, Methodology, Investigation, Formal analysis, Validation, Writing – Original Draft

**Stine Møllerud Fransen**: Conceptualization, Methodology, Investigation

**Darryl S. Pickering**: Conceptualization, Methodology, Investigation, Formal analysis, Validation, Writing – Review & Editing, Supervision

**Karla Frydenvang**: Investigation, Formal analysis, Validation, Writing – Review & Editing, Supervision

**Pierre Francotte**: Resources, Writing – Review & Editing

**Bernard Pirotte**: Resources, Writing – Review & Editing

**Anders Skov Kristensen**: Methodology, Validation, Writing – Review & Editing, Supervision

**Jette Sandholm Kastrup**: Conceptualization, Methodology, Formal analysis, Validation, Writing – Original Draft, Review & Editing, Supervision, Funding acquisition

## Declarations of interest

We declare no competing interests.

## PDB references

The structure coordinates and corresponding structure factor file of the GluK1-LBD dimer in complex with kainate and BPAM538 have been deposited in the Protein Data Bank under the accession code XXXX.

## Abbreviations

AMPA: α-amino-3-hydroxy-5-methylisoxazole-4-propionate
GluK1-LBD: ligand-binding domain of GluK1
iGluRs: ionotropic glutamate receptors
LBD: ligand-binding domain
RFU: relative fluorescence unit

## Notes

### Competing Interest Statement

The authors have declared no competing interest.

